# Distinct properties of layer 3 pyramidal neurons from prefrontal and parietal areas of the monkey neocortex

**DOI:** 10.1101/649228

**Authors:** Guillermo Gonzalez-Burgos, Takeaki Miyamae, Yosef Krimer, Yelena Gulchina, Diego Pafundo, Olga Krimer, Holly Bazmi, Dominique Arion, John F Enwright, Kenneth Fish, David A Lewis

**Affiliations:** Translational Neuroscience Program, Department of Psychiatry, University of Pittsburgh School of Medicine. W1647, Biomedical Science Tower, 200 Lothrop Street, Pittsburgh PA 15261

## Abstract

In primates, working memory function depends on activity in a distributed network of cortical areas that display different patterns of delay task-related activity. These differences are correlated with, and might depend on, distinctive properties of the neurons located in each area. For example, layer 3 pyramidal neurons (L3PNs) differ significantly between primary visual and dorsolateral prefrontal (DLPFC) cortices. However, to what extent L3PNs differ between DLPFC and other association cortical areas is less clear. Hence, we compared the properties of L3PNs in monkey DLPFC versus posterior parietal cortex (PPC), a key node in the cortical working memory network. Using patch clamp recordings and biocytin cell filling in acute brain slices, we assessed the physiology and morphology of L3PNs from monkey DLPFC and PPC. The L3PN transcriptome was studied using laser microdissection combined with DNA microarray or quantitative PCR. We found that in both DLPFC and PPC, L3PNs were divided into regular spiking (RS-L3PNs) and bursting (B-L3PNs) physiological subtypes. Whereas regional differences in single-cell excitability were modest, B-L3PNs were rare in PPC (RS-L3PN:B-L3PN, 94:6), but were abundant in DLPFC (50:50), showing greater physiological diversity. Moreover, DLPFC L3PNs display larger and more complex basal dendrites with higher dendritic spine density. Additionally, we found differential expression of hundreds of genes, suggesting a transcriptional basis for the differences in L3PN phenotype between DLPFC and PPC. These data show that the previously observed differences between DLPFC and PPC neuron activity during working memory tasks are associated with diversity in the cellular/molecular properties of L3PNs.

**Significance statement:** In the human and non-human primate neocortex, layer 3 pyramidal neurons (L3PNs) differ significantly between dorsolateral prefrontal (DLPFC) and sensory areas. Hence, L3PN properties reflect, and may contribute to, a greater complexity of computations performed in DLPFC. However, across association cortical areas, L3PN properties are largely unexplored. We studied the physiology, dendrite morphology and transcriptome of L3PNs from macaque monkey DLPFC and posterior parietal cortex (PPC), two key nodes in the cortical working memory network. L3PNs from DLPFC had greater diversity of physiological properties and larger basal dendrites with higher spine density. Moreover, transcriptome analysis suggested a molecular basis for the differences in the physiological and morphological phenotypes of L3PNs from DLPFC and PPC.

## Introduction

In primates, working memory function depends on activity in a distributed network of cortical areas (Leavitt et al., 2017; Dotson et al., 2018). Within this network, sensory and association areas display different activity patterns during working memory tasks, reflecting the transformation of sensory input into a behavioral response across a delay (Christophel et al., 2017). For example, neurons in primary visual cortex (V1) increase their firing rate during stimulus presentation, whereas neurons in the dorsolateral prefrontal cortex (DLPFC) often change their activity during the delay period (Dotson et al., 2018; Yang et al., 2018; Quentin et al., 2019). These regional differences might reflect the hierarchy of activity timescales observed in the primate neocortex (Murray et al., 2014).

The hierarchy of activity across the working memory network might depend, at least in part, on distinctive properties of the neurons located in each area (Murray et al., 2014). Consistent with this idea, layer 3 pyramidal neurons (L3PNs) in monkey V1 are more excitable, than L3PNs in DLPFC (Amatrudo et al., 2012; Gilman et al., 2017). Moreover, L3PNs from monkey DLPFC have larger dendrites with higher dendritic spine densities than L3PNs from V1 (Elston, 2000, 2003; Medalla and Luebke, 2015). Furthermore, in both monkeys and humans, the gene expression profile, a major determinant of L3PN physiology and morphology (Cubelos et al., 2010; Tripathy et al., 2017), differs markedly in layer 3 between V1 and DLPFC (Bernard et al., 2012; Hoftman et al., 2018; Zhu et al., 2018). These findings suggest that the physiology of L3PNs differs across cortical areas in association with differences in their dendritic tree properties and transcriptome.

The functional differences between monkey V1 and DLPFC during working memory tasks might be the product, at least in part, of regional differences in the properties of L3PNs that provide a cellular substrate for the complex computations performed in DLPFC (Miller and Cohen, 2001). However, the extent to which L3PNs differ between DLPFC and other association cortical areas in the working memory network is less clear, as the limited data currently available suggest both similarities and differences. For example, neurons in the primate posterior parietal cortex (PPC), a key node in the working memory network, also exhibit delay-related activity during working memory tasks (Constantinidis and Steinmetz, 1996; Chafee and Goldman-Rakic, 1998), and PPC inactivation affects DLPFC neuron activity and working memory task performance (Quintana et al., 1989; Chafee and Goldman-Rakic, 2000).

Moreover, the PPC shares many cytoarchitectonic features (Beul et al., 2017; Goulas et al., 2018) and patterns of cortico-cortical connectivity (Selemon and Goldman-Rakic, 1988; Markov et al., 2014) with the DLPFC, and is interconnected with DLPFC via projections from L3PNs (Andersen et al., 1990; Medalla and Barbas, 2006; Markov et al., 2014). These and other findings suggest that the PPC and DLPFC constitute similar network hubs, and perform similar high level computations (Freedman and Ibos, 2018), suggesting that, between PPC and DLPFC, L3PNs have comparable properties. However, PPC and DLPFC neurons are separated in the hierarchy of activity timescales in the primate cortex (Murray et al., 2014). In addition, the working memory task-related activity of PPC neurons is less generalized (Sarma et al., 2016), may encode different aspects of task-related information (Qi et al., 2015), and is more sensitive to the effects of distractors (Constantinidis and Steinmetz, 1996). Moreover, gene expression differs markedly between frontal and parietal regions of the primate neocortex (Bernard et al., 2012; Hoftman et al., 2018; Zhu et al., 2018).

Here, we assessed the physiology, morphology, and transcriptome of L3PNs in monkey DLPFC and PPC. We found that despite modest regional differences in single-cell excitability, L3PNs had greater physiological diversity and morphological complexity in the DLPFC than in the PPC. Moreover, the differential expression of hundreds of genes suggested a molecular/transcriptional basis for the differences in L3PN phenotype between DLPFC and PPC.

## Materials and methods

### Animals

All housing and experimental procedures were conducted in accordance with USDA and NIH guidelines and were approved by the University of Pittsburgh Institutional Animal Care and Use Committee. For the electrophysiology experiments, tissue was obtained from 9 rhesus monkeys (*Macaca mulatta*, 7 males and 2 females; 27 to 47 months of age) that were experimentally naïve until entry into this study. Tissue sections from 5 of these monkeys were used for microarray analyses of gene expression in L3PNs. An independent cohort of 7 male rhesus macaque monkeys (41-44 months of age), were used for quantitative PCR (qPCR) studies of L3PNs. These animals served as the vehicle-exposed group in studies investigating the impact of Δ^9^-tetrahydrocannabinol on cognitive task performance (Verrico et al., 2014).

### Brain slice preparation

Tissue blocks (Figure 1A) containing both banks of the principal sulcus (DLPFC area 46) or the intraparietal sulcus and adjacent lateral cortex (PPC areas LIP and 7a) were obtained after the animals were deeply anesthetized and perfused transcardially (Gonzalez-Burgos et al., 2015) with ice-cold sucrose-modified artificial cerebro-spinal fluid (sucrose-ACSF, in mM): sucrose 200, NaCl 15, KCl 1.9, Na_2_HPO_4_ 1.2, NaHCO_3_ 33, MgCl_2_ 6, CaCl_2_ 0.5, glucose 10 and kynurenic acid 2; pH 7.3–7.4 when bubbled with 95% O_2_-5% CO_2_. From one of the female monkeys, tissue slices were prepared from both DLPFC and PPC areas. For all other experiments, slices were obtained from either DLPFC (4 male monkeys) or PPC (3 males, 1 female). Slices were cut in the coronal plane at 300 μm thickness, in a vibrating microtome (VT1000S, Leica Microsystems) while submerged in ice-cold sucrose-ACSF. Immediately after cutting, the slices were transferred to an incubation chamber filled with the following room temperature ACSF (mM): NaCl 125, KCl 2.5, Na_2_HPO_4_ 1.25, glucose 10, NaHCO_3_ 25, MgCl_2_ 1 and CaCl2 2, pH 7.3–7.4 when bubbled with 95% O_2_-5% CO_2_. Electrophysiological recordings were initiated 1 to 14 hours after tissue slicing was completed. All chemical reagents used to prepare solutions were purchased from Sigma Chemicals Company.

**Figure 1.**
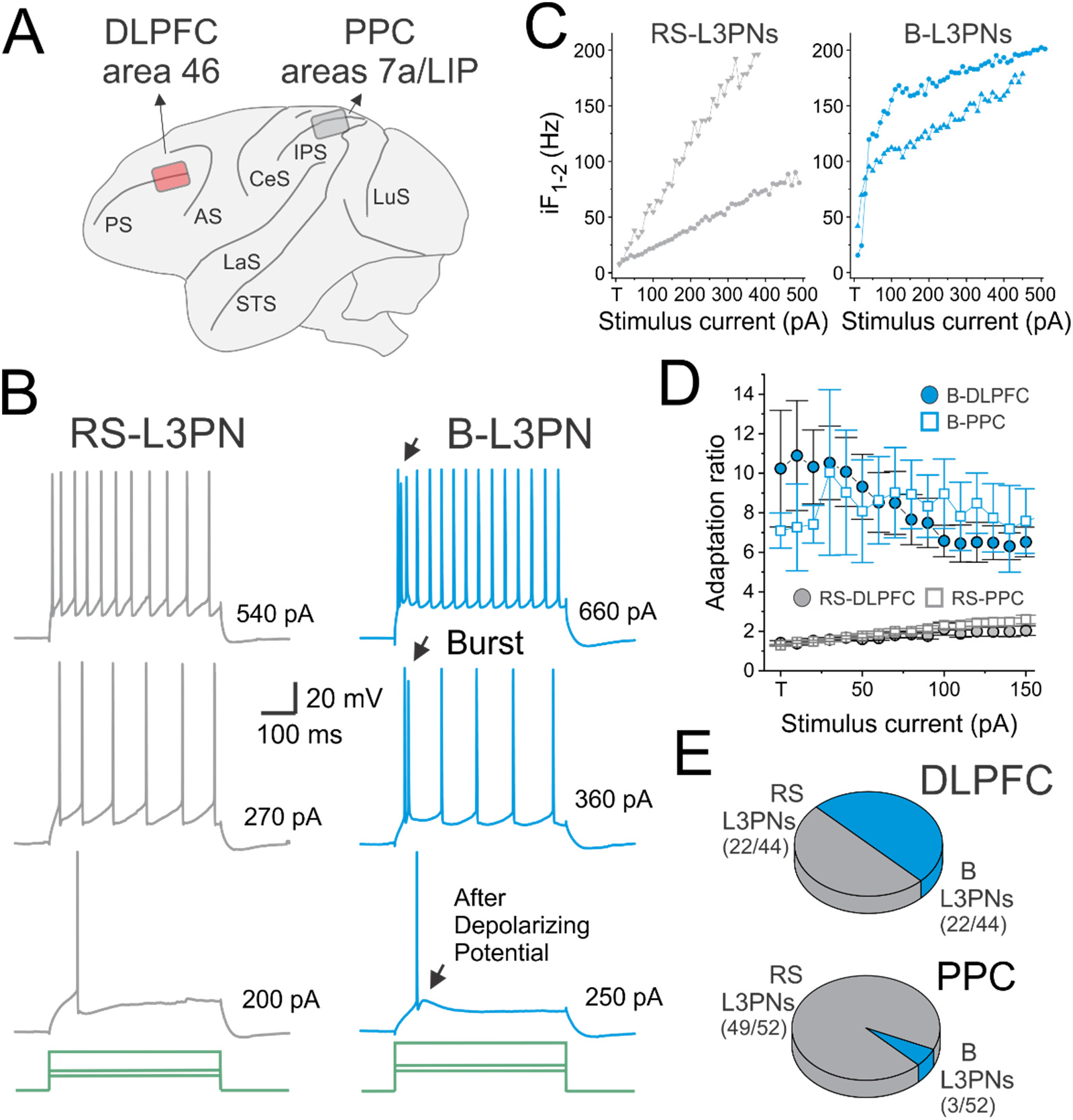
Firing patterns of L3PNs recorded in slices from monkey DLPFC and PPC. **A)** Schematic lateral view of the macaque monkey cortex, displaying the approximate location of tissue blocks obtained to prepare slices from DLPFC area 46 or from PPC areas 7a and LIP. PS: principal sulcus; AS: arcuate sulcus; CeS: central sulcus; LaS: lateral sulcus; IPS: intraparietal sulcus; STS: superior temporal sulcus; LuS: lunate sulcus. **B)** Examples of firing patterns of L3PNs studied ex vivo using current clamp recordings. The left panel show an example RS-L3PN recorded from PPC, and the right panel illustrates recordings from a B-L3PN in a DLPFC slice. Note the presence of an after depolarizing potential in the B-L3PN, which eventually triggers a burst of 2-3 action potentials at the onset of the response. **C)** Plots of the instantaneous frequency for the first two action potentials (iF_1-2_) fired in response to excitatory current injection. The left plots are two examples of RS-L3PNs, and the right plots are two examples of B-L3PNs. Note that RS-L3PNs reach iF_1-2_ >= 100 Hz only at input currents 100 pA or more above current threshold, whereas B-L3PNs reach 100 Hz typically within 50 pA of current threshold. **D)** Plots illustrating the differences in the spike frequency adaptation between RS-L3PNs and B-L3PNs. Adaptation was quantified dividing the last inter-spike interval by the first, and this ratio was plotted as a function of stimulus current above threshold. **E)** Pie charts illustrating the significantly different proportions of RS-L3PNs and B-L3PNs in DLPFC and PPC. The difference in proportions of RS-L3PNs and B-L3PNs was highly significant (p=6.9×10^-7^, Chi Square test).

### Electrophysiological recordings

Slices were placed in a recording chamber superfused at 2-3 ml/min with the following ACSF (mM): NaCl 125; KCl 2.5; Na_2_HPO_4_ 1.25; NaHCO_3_ 25; glucose 10; CaCl_2_ 2; MgCl_2_ 1; CNQX 0.01, bubbled with 95 % O_2_ / 5 % CO_2_ at 30-32 °C. Whole-cell recordings were obtained from L3PNs identified visually by infrared differential interference contrast video microcoscopyusing Olympus or Zeiss microscopes equipped with CCD video cameras (EXi Aqua, Q-Imaging). Recordings were obtained from L3PNs located in either medial or lateral banks of the principal sulcus in DLPFC area 46, or in cytoarchitectonic areas LIP and 7a of the PPC, in the lateral bank of the intraparietal sulcus. Recording pipettes had 3-5 MΩ resistance when filled with the following solution (mM): KGluconate 60; KCl 70; NaCl 10; EGTA 0.2; HEPES 10; MgATP 4; NaGTP 0.3, NaPhosphocreatine 14, biocytin 0.4 %, pH 7.2-7.3, adjusted with KOH. Current clamp recordings and data analysis were conducted as described previously (Henze et al., 2000; Gonzalez-Burgos et al., 2004; Zaitsev et al., 2012), with Multiclamp 700A or 700B amplifiers (Axon Instruments) operating in bridge mode with pipette capacitance compensation. Recordings were included in data analysis only if the resting membrane potential was ≤ −60 mV.

#### Pyramidal cell membrane properties

To measure membrane properties, families of 500 ms current steps were used (−80 to 600 pA, incrementing by 10 pA, 2-3 repeats per current level). The input resistance was estimated via the voltage response to current injection in the −50 to −10 pA range, which was well-fit by a linear relation in each L3PN. The membrane time constant was estimated from the voltage response averaged for hyperpolarizing current steps of −30 to −10 pA amplitude, using single exponential functions fit to the voltage response. This measure is an approximation of the actual membrane time constant, which, in a passive neuron with complex geometry, is the time constant of the slowest component of a multiexponential time course (Spruston et al., 1994). The percentage of sag was determined by estimating the hyperpolarization level during the last 10 ms of the response to the −50 pA current step, as a percentage of the hyperpolarization measured 20 ms after the onset of the hyperpolarizing current step, during a 10 ms window. The current threshold, or rheobase, was defined as the first current level producing spikes in at least 2 of the 3 repeats for that current level. The voltage threshold to fire an action potential (AP threshold) was defined based on derivatives of the membrane potential (Henze et al. 2000). The afterhyperpolarization (AHP) amplitude was defined as the voltage from AP threshold to the trough of the AHP. The AP width, or duration, at half maximal amplitude (AP HW) was defined as the duration of the AP at 50% of its maximal amplitude. All AP parameters were estimated from single APs (2-3 repeats per neuron), and measured on the first AP fired in response to rheobase current stimulation. The slope of the relation between mean firing frequency and current step amplitude (f–I plot) was estimated from the linear region of the plots of mean firing frequency versus stimulus amplitude. The mean firing frequency was calculated from the number of APs evoked per 500 ms stimulus, averaged for the 2-3 repetitions of each stimulus amplitude. The adaptation ratio was measured as the ratio between the last and first inter-spike intervals. To classify each neuron as bursting or regular firing, we measured the instantaneous firing frequency from the first two APs evoked by current injection at various current levels above rheobase (Figure 1C).

### Histological processing and morphological reconstruction of biocytin-filled neurons

During recordings, L3PNs were filled with 0.5% biocytin, and then visualized and reconstructed using procedures described previously (Zaitsev et al., 2012). Briefly, after recordings, the slices were immersed in 4% p-formaldehyde in 0.1 M phosphate-buffered saline (PBS) for 24-72 h at 4 °C. The slices were stored at −80 °C in a cryo-protection solution (33% glycerol, 33% ethylene glycol, in 0.1 M PBS) until processed. To visualize biocytin, the slices were resectioned at 60 μm, incubated with 1% H_2_O_2_, and immersed in blocking serum containing 0.5% Triton X-100 for 2–3 h at room temperature. The tissue was then rinsed and incubated with the avidin–biotin–peroxidase complex (1:100; Vector Laboratories) in PBS for 4 h at room temperature. Sections were rinsed, stained with the Nickel-enhanced 3,3’-diaminobenzidine chromogen, mounted on gelatin-coated glass slides, dehydrated, and cover slipped. Three-dimensional reconstructions of the dendritic arbor were performed using the Neurolucida tracing system (MBF Bioscience). Dendritic spines were identified using differential interference contrast images of the biocytin-filled dendrites (Figure 5). Although multiple spine morphologies were found, including stubby, mushroom, and thin spines (Figure 5), morphological subtypes of spines were not distinguished in the analysis of basal dendrite spine density. For the measurements of spine density, a single primary basal dendrite was randomly selected for each L3PN, and spines were reconstructed throughout the entire length of the basal dendrite. The peak spine density was defined as the highest spine number per micron value observed in each basal dendrite analyzed. The mean spine density was the average of the spine number per micron values obtained for each basal dendrite analyzed.

### Statistical analysis of electrophysiological and morphological data

The data were expressed as means ± S.E.M, except when otherwise indicated. To assess normality of the distribution of the data we used the Shapiro-Wilk test applied to the residuals of the data. The Shapiro-Wilk tests rejected normal distribution for 5 of the electrophysiological parameters measured (Input resistance, p=0.00000152; Rheobase, p=0.000168; Membrane time constant, p=5.6 × 10^-13; Slope of f-I plot, p=0.00202; AP HW, p=0.0221). For the data on dendritic tree morphology and dendritic spine density, normal distribution was rejected for 2 of the measured parameters (Oblique apical dendrite length, p=0.0357; Apical dendrite tuft length, p=0.0063). For the cases in which normality of the distribution was rejected, the Shapiro-Wilk tests were repeated after natural logarithm transformation of the data. Normality of the distribution was still rejected for the log-transformed of two of the electrophysiological parameters (Rheobase, p=0.0109; Membrane time constant, p=0.00146), and for the Oblique apical dendrite length (p=0.0127). When normality of the distribution of the log-transformed data was rejected, we employed a non-parametric test, as indicated in each case. Group means were compared using Student’s t-test, One-Way ANOVA, Mann-Whitney’s U test, Kruskal-Wallis ANOVA, or Chi Squared test, as indicated in each case. Statistical tests were performed using SPSS (IBM Corp.).

### Transcriptome analysis

#### Laser microdissection

Coronal cryostat sections (12 μm thick) containing the principal sulcus or the intraparietal sulcus were cut from each monkey, mounted onto polyethylene naphthalate (PEN) membrane slides (Leica Microsystems, Bannockburn, IL, USA) and stained for Nissl substance using a rapid procedure as previously described (Datta et al., 2015). L3PNs were identified based on their characteristic somal morphology and the presence of a prominent apical dendrite directed radially toward the pia surface. From each monkey, individual L3PNs were dissected from the dorsal and ventral banks of the principal sulcus (DLPFC area 46) or from the lateral bank of the intraparietal sulcus (PPC areas LIP and 7a) and pooled into samples of 100 or 500 cells for microarray profiling or qPCR, respectively.

#### DNA microarray profiling

RNA was extracted from a pool of 100 L3PNs using the RNeasy® Plus Micro Kit (QIAGEN) and subjected to a single round of amplification using the Ovation Pico WTA system, labeled using the Encore Biotin module (Nugen) and loaded for transcriptome analysis on a GeneChip™ Rhesus Macaque Genome Array (ThermoFisher scientific), designed to assess expression levels of transcripts in the macaque monkey genome. For each sample, expression intensities were extracted from Affymetrix Expression Console using the RMA method (Irizarry et al., 2003) and transformed to log-scale (base 2). The microarray analysis was performed consecutively in three separate batches, and initial principal component analysis identified a strong influence of processing batch on transcript levels. Therefore, the Combat function of the R SVA package (Leek et al., 2018) was used to correct for batch effect. Any probe in which all samples had an expression level below 4 was considered to represent background and was removed, resulting in 23,748 probes for differential expression analysis. All replicate samples from a given region within an animal (2-3 replicates per animal) were averaged and then mean expression across all 5 animals were calculated for each region. Differential expression between regions was determined using a paired Student’s t-test followed by correction for multiple comparisons using Storey’s method (Storey, 2002).

#### Quantitative PCR

For qPCR verification, total RNA was extracted from pools of 500 L3PNs from each region and converted to complementary DNA using the Superscript IV VILO Mastermix (ThermoFisher). The PCR amplification was performed using Power SYBR green dye and ViiA 7 Real-Time PCR system (Applied Biosystems, Carlsbad, CA). All primer sets used in the qPCR analysis (Table 3) had a minimum primer efficiency of 97%, and all amplified products resulted in a specific single product in dissociation curve analysis. β-Actin (ACTB), guanine nucleotide-binding protein G(s) subunit alpha (GNAS), and glyceraldehyde-3-phosphate dehydrogenase (GAPDH), were used as endogenous reference genes. The use of these normalizer genes was supported by the microarray data showing an absence of difference in expression levels of these genes between DLPFC and PPC samples. The difference in cycle thresholds (dCTs) was calculated for each sample by subtracting the geometric mean of the 3 normalizers from the CT of the target transcript. Since dCT represents the log2-transformed expression ratio of each target transcript to the reference genes, the relative level of the target transcript for each subject is reported as 2^-dCT^.

#### Validation of cell type-specificity of laser microdissection

To confirm that L3PN samples collected by laser microdissection were enriched in PN markers, we assessed transcripts levels of *SLC17A7*, the gene encoding the vesicular glutamate transporter 1 (a glutamatergic PN marker), and *SLC32A1*, the gene encoding the vesicular GABA transporter 1 (a GABAergic neuron marker). We found an enrichment of SLC17A7 to SLC32A1 that was on average 268-fold in DLPFC and 143-fold in PPC samples, respectively, confirming the enrichment in PN transcripts.

## Results

### DLPFC L3PNs display greater electrophysiological diversity

We assessed the action potential firing pattern and other intrinsic membrane properties in current clamp recordings obtained from L3PNs in slices from DLPFC area 46 or PPC areas 7a and LIP (Figure 1A). The response of the L3PNs to excitatory current injection revealed two distinct firing pattern subtypes (Figure 1B), here termed regular spiking (RS-L3PNs) and bursting (B-L3PNs). RS-L3PNs exhibited progressive and relatively weak adaptation of the firing frequency during current injection. B-L3PNs displayed a transient depolarizing potential following the first action potential, which triggered a burst of action potentials near the onset of the stimulus and was followed by substantial spike frequency adaptation. Individual L3PNs were classified as RS-L3PN or B-L3PN by plotting the instantaneous firing frequency for the first two action potentials (iF_1-2_) versus stimulus current (Figure 1C). In RS-L3PNs, the iF_1-2_ increased progressively with stimulus amplitude, reaching ~100 Hz frequency values at stimulus currents ≥ 200 pA above rheobase (Figure 1C). In contrast, the iF_1-2_ increased steeply in B-L3PNs, reaching 100 Hz within 50 pA of rheobase (Figure 1C), as previously reported for pyramidal neurons in rodent cortex (Graves et al., 2012). While showing a steeper increase of iF_1-2_ as a function of input current (Figure 1C), B-L3PNs also had stronger firing frequency adaptation than RS-L3PNs (Figure 1D), as previously reported for monkey DLPFC L3PNs (Zaitsev et al., 2012). Spike frequency adaptation, however, did not differ between cortical areas for each type of L3PN (Figure 1D).

In DLPFC, the recorded neurons were equally divided between RS-L3PNs (22/44) and B-L3PNs (22/44) (Figure 1E), whereas in PPC, 94.2% (49/52) of the recorded neurons were RS-L3PNs and only 5.8% were B-L3PNs (Figure 1E). This regional difference in proportions of the two cell types was highly significant (p=6.9×10^-7^, Chi Square test).

To test for differences in intrinsic membrane properties, we compared various electrophysiological parameters of L3PNs from DLPFC and PPC. Given the few B-L3PNs found in PPC (n=3), the differences between physiological subtypes of L3PNs were tested by comparing B-L3PNs versus RS-L3PNs in the DLPFC, and the differences between DLPFC and PPC were tested comparing RS-L3PNs from each region. Of the physiological parameters assessed (Figure 2), four differed significantly between DLPFC B-L3PNs and RS-L3PNs, as previously reported (Zaitsev et al., 2012). These differences include less spike frequency adaptation (Figure 1), higher input resistance, smaller hyperpolarizing response sag, and larger afterhyperpolarization amplitude in RS-L3PNs (Figure 2). Higher input resistance and smaller sag may both reflect lower density of hyperpolarization-activated channels, which mediate a current that decreases membrane excitability, suggesting neurons with smaller sag are more excitable. A larger AHP amplitude may help resetting the voltage threshold during the interval between subsequent spikes. These differences suggest that, despite the absence of difference in action potential threshold, resting membrane potential and other parameters, RS-L3PNs might be more excitable than B-L3PNs, and hence more likely to be recruited by similar levels of excitatory input.

**Figure 2.**
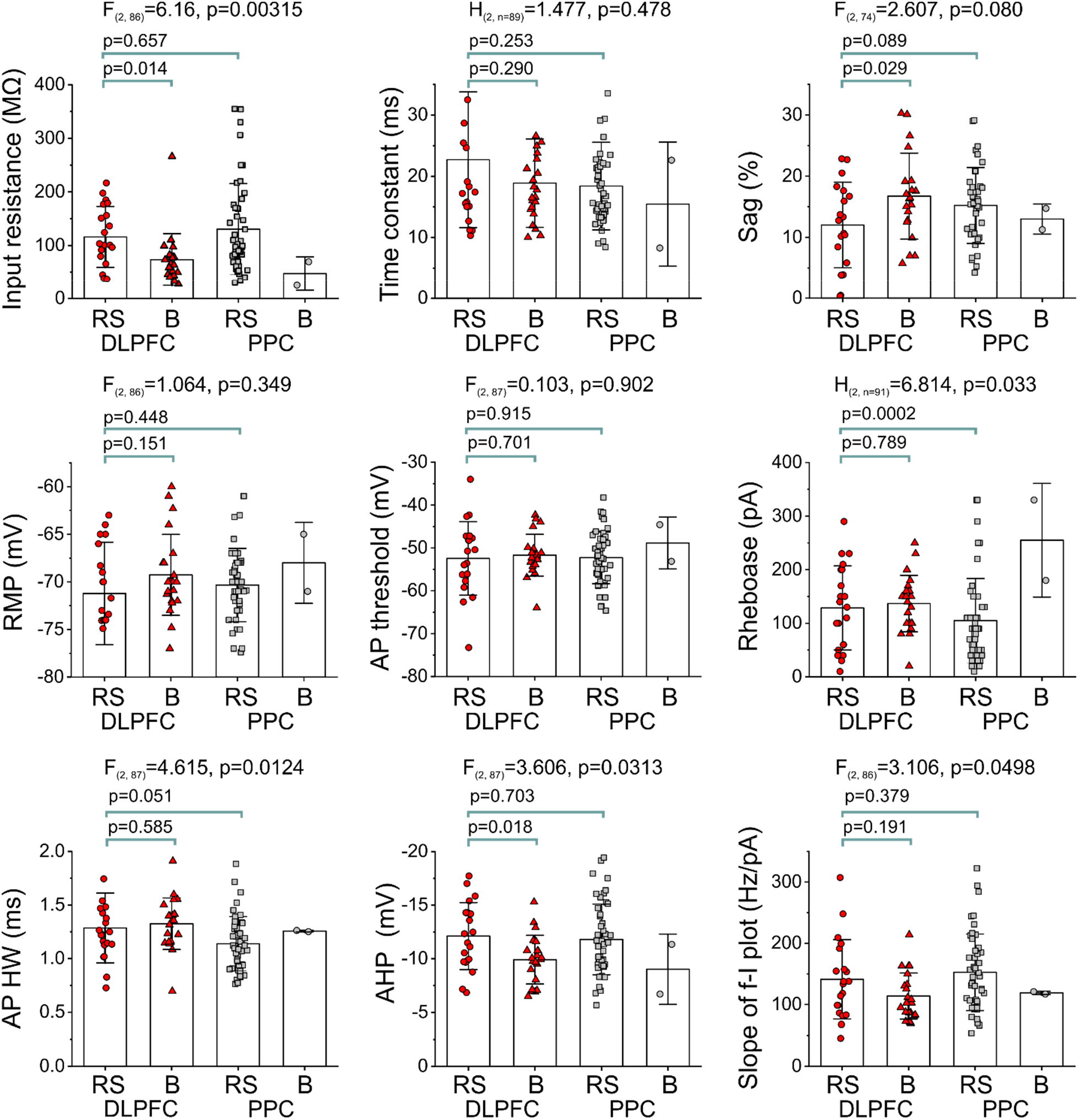
Intrinsic physiological properties of L3PNs assessed in current clamp recordings in DLPFC or PPC brain slices. The recorded neurons were classified as RS-L3PNs (here abbreviated as RS) or B-L3PNs (here abbreviated as B), using iF_1-2_ versus current plots as illustrated in Figure 1C. The membrane properties reported in the figure were measured as described in the methods section. Shown are the results of One-Way ANOVA analysis, or non-parametric Friedman’s ANOVA followed by post-hoc contrasts. For two parameters (membrane time constant and Rheobase) normality tests showed significant deviation from the normal distribution of the raw or log-transformed data, hence the non-parametric test was employed. As the number of B-L3PNs found in PPC slices was small (n=3), and membrane properties could be characterized in just 2 of these 3 B-L3PNs, these were not included in the statistical analysis. RMP: resting membrane potential. AP threshold: voltage threshold for firing an action potential. AP HW: duration, or width, of the action potential at half maximal amplitude. AHP: amplitude of the after hyperpolarizing potential. Slope of f-I plot: slope of the linear portion of the relation between mean firing frequency and stimulus current.

Bursting depends on the activity of T-type voltage-gated Ca^2+^ channels that show pronounced inactivation at relatively depolarized steady-state membrane resting potentials (Williams and Stuart, 1999; Clarkson et al., 2017; Dumenieu et al., 2018). Hence, a more depolarized resting membrane could inactivate the T-type channels mediating burst firing. However, the resting membrane potential did not differ between DLPFC B-L3PNs and RS-L3PNs (Figure 2) from either DLPFC (p=0.151) or PPC (p=0.448), indicating that the presence or absence of bursting was not due to differences in steady-state inactivation of T-type channels between samples. Whereas 4 of 10 parameters assessed differed between RS-L3PNs and B-L3PNs, only the threshold current, or rheobase, differed significantly between RS-L3PNs from DLPFC and PPC, being slightly lower in PPC L3PNs (Figure 2). Hence, our data suggest modest differences in the membrane properties and single-cell excitability between DLPFC and PPC L3PNs.

### DLPFC L3PNs have larger and more complex basal dendritic trees with higher spine density

Dendritic tree morphology was reconstructed (Figure 3) for 21 DLPFC and 18 PPC L3PNs filled with biocytin during the electrophysiological recordings. The distance from the pial surface to the cell bodies (i.e., soma depth) varied among the reconstructed L3PNs (Figure 3A,B), ranging from 329 to 655 μm in the DLPFC, and from 270 to 714 μm in the PPC, well within the boundaries of layer 3 as measured in Nissl-stained sections. The mean soma depth of DLPFC (491 ± 23 μm) and PPC (506 ± 27 μm) L3PNs did not differ (t=0.456, p=0.651) between areas. The results of the quantitative morphometric analysis are depicted in Figure 4.

**Figure 3.**
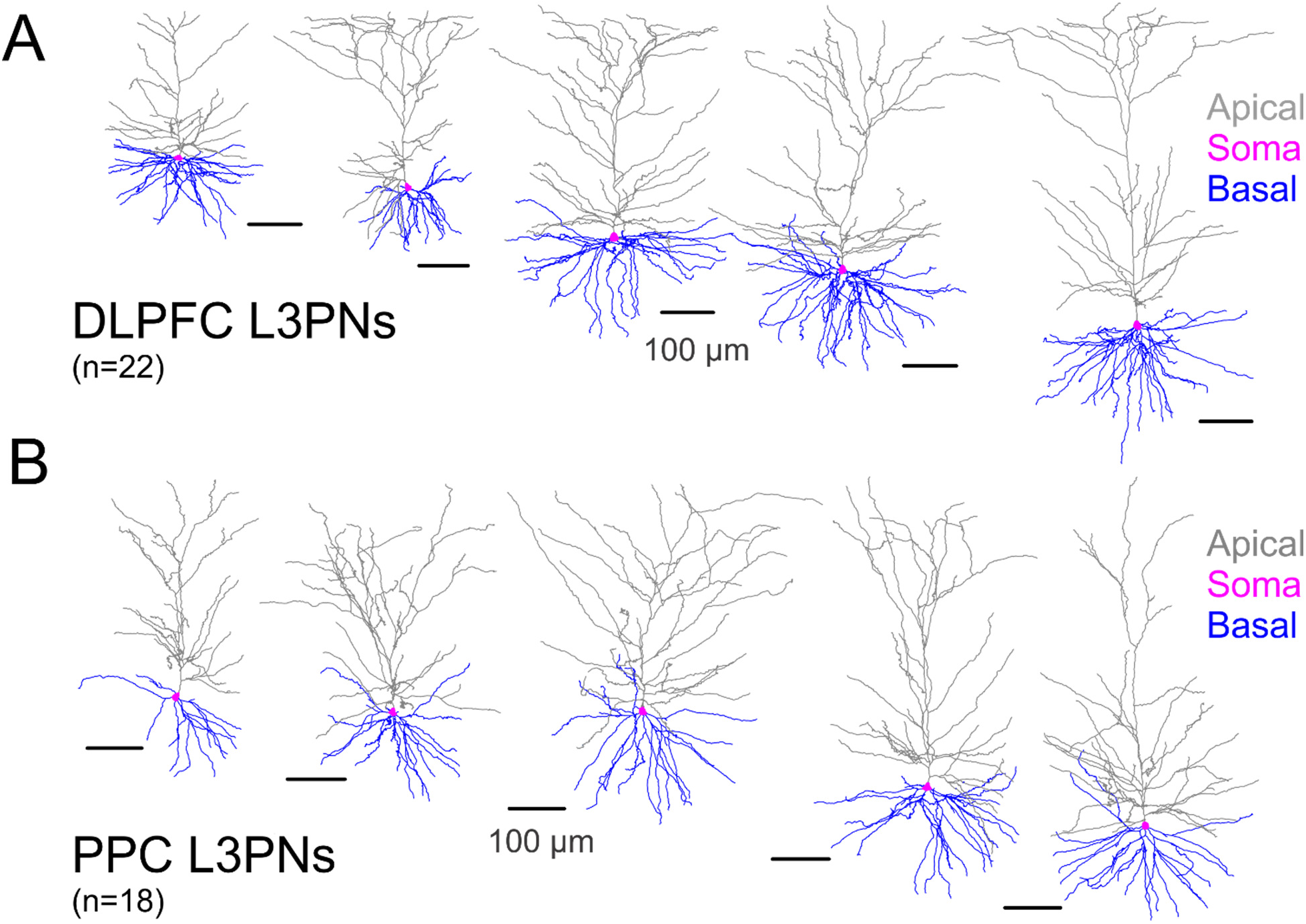
Two-dimensional views of the three-dimensional reconstructions of the apical and basal dendritic trees of DLPFC and PPC L3PNs. **A)** Examples of reconstructions of the apical (grey) and basal (blue) dendrites of L3PNs from DLPFC. The soma is depicted in magenta color. **B)** Examples of reconstructions of the apical and basal dendrites of L3PNs from PPC. Calibration bars apply to both A and B.

**Figure 4.**
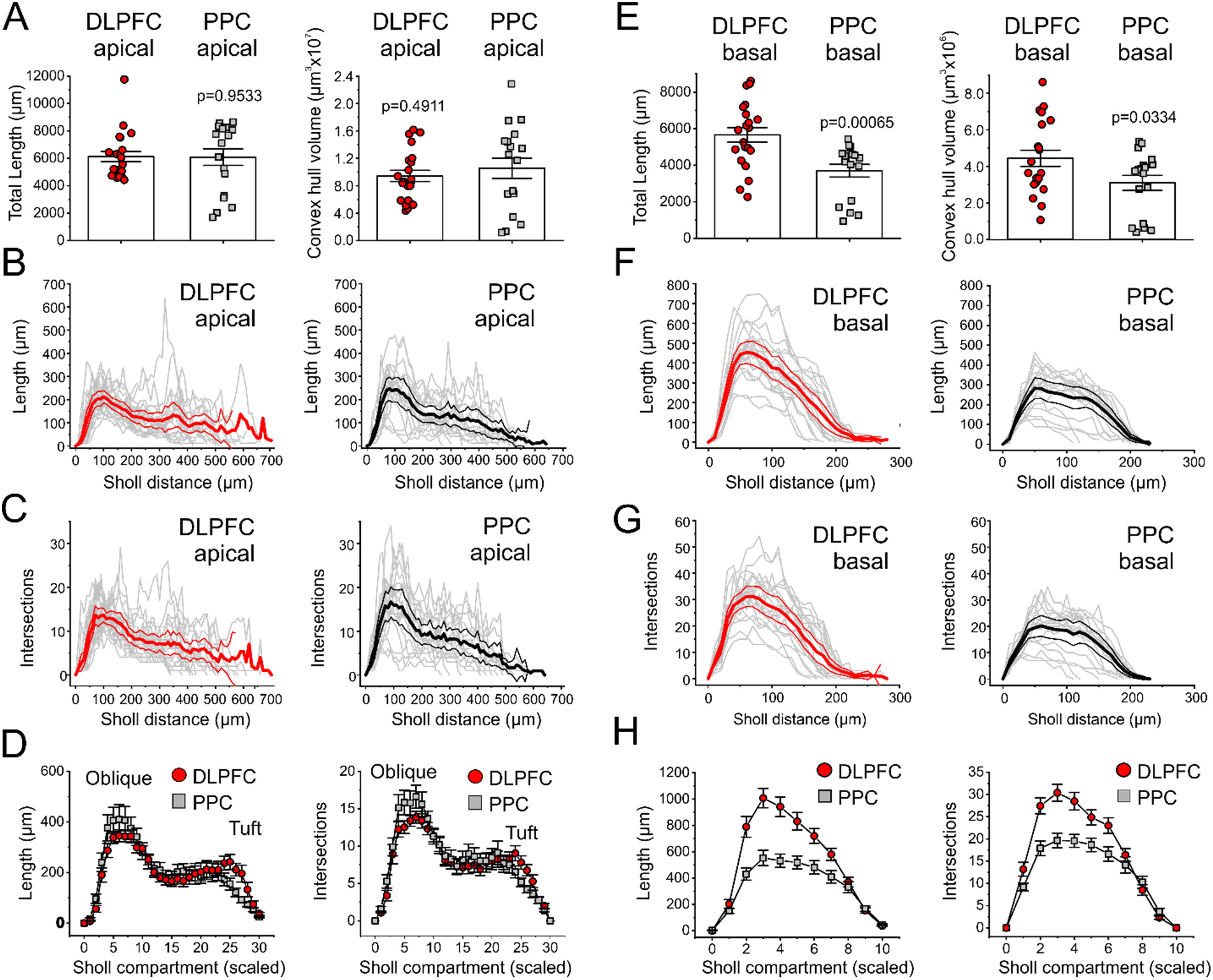
Quantitative analysis of apical dendrite properties of DLPFC and PPC L3PNs. **A)** Left: Bar graphs illustrating the total length of the apical dendrites for DLPFC (n=21) and PPC (n=18) L3PNs. Shown are the results of Student’s t-test comparisons. Right: Bar graphs illustrating the convex hull volume of the apical dendrites for the same DLPFC and PPC L3PNs. Shown are the results of a Student’s t-test comparison. **B)** Sholl analysis of apical dendrites: Plots of dendrite length as a function of Sholl distance from the soma for the apical dendrites of DLPFC (left) and PPC (right) L3PNs. The thick lines indicate the mean value and the dark lines above and below are the 95% confidence intervals. The gray lines represent data from individual L3PNs. **C)** Sholl analysis of apical dendrites: Plots of number of intersections as a function of Sholl distance from the soma for the apical dendrites of DLPFC (left) and PPC (right) L3PNs. The different lines represent the data as indicated for C). **D)** Sholl analysis of apical dendrites: Scaled Sholl plots obtained for dendrite length (left) and number of intersections (right) after the apical dendrites of each L3PN were divided into 30 compartments, by adjusting the increment of Sholl radius for each neuron. For statistical comparisons, see Table 3. **E)** Left: Bar graphs illustrating the total length of the basal dendrites for DLPFC (n=21) and PPC (n=18) L3PNs. Shown are the results of a Student’s t-test comparisons. Right: Bar graphs illustrating the convex hull volume of the basal dendrites for the same DLPFC and PPC L3PNs as in A). Shown are the results of a Student’s t-test comparison. **F)** Sholl analysis of basal dendrites: Plots of dendrite length as a function of Sholl distance from the soma for the basal dendrites of DLPFC (left) and PPC (right) L3PNs. The thick lines indicate the mean value and the dark lines above and below are the 95% confidence intervals. The gray lines represent data from individual L3PNs. **G)** Sholl analysis of basal dendrites: Plots of number of intersections as a function of Sholl distance from the soma for the basal dendrites of DLPFC (left) and PPC (right) L3PNs. The different lines represent the data as indicated in C). **H)** Sholl analysis of basal dendrites: Scaled Sholl plots obtained for dendrite length (left) and number of intersections (right) after the basal dendrites of each L3PN were divided into 10 compartments, by adjusting the increment of Sholl radius for each neuron. For statistical comparisons, see Table 3.

Quantitative analyses of the apical dendrites (Figure 4A) revealed that neither the total length nor the convex hull volume, a three-dimensional parameter estimating the tissue volume where the dendrites sample inputs, differed between DLPFC and PPC L3PNs. Sholl analysis performed to obtain measures of length (Figure 4B) and complexity (Figure 4C) across compartments of the apical dendrites, likewise suggested an absence of difference between DLPFC and PPC L3PNs. However, the reconstructed L3PNs had different soma depths (Figure 3), and therefore, in the Sholl plots, proximal dendrite compartments of some neurons were aligned with more distal dendrites of other cells. To reduce this misalignment, we built scaled Sholl plots dividing the apical dendrites of all L3PNs into an equal number of compartments.

The plots for scaled apical dendrites largely overlapped between DLPFC and PPC L3PNs (Figure 4D), and neither the peak length nor the peak number of intersections differed significantly between cortical areas in the oblique branch region or tuft of the apical dendrites (Table 2).

**Table 1.**
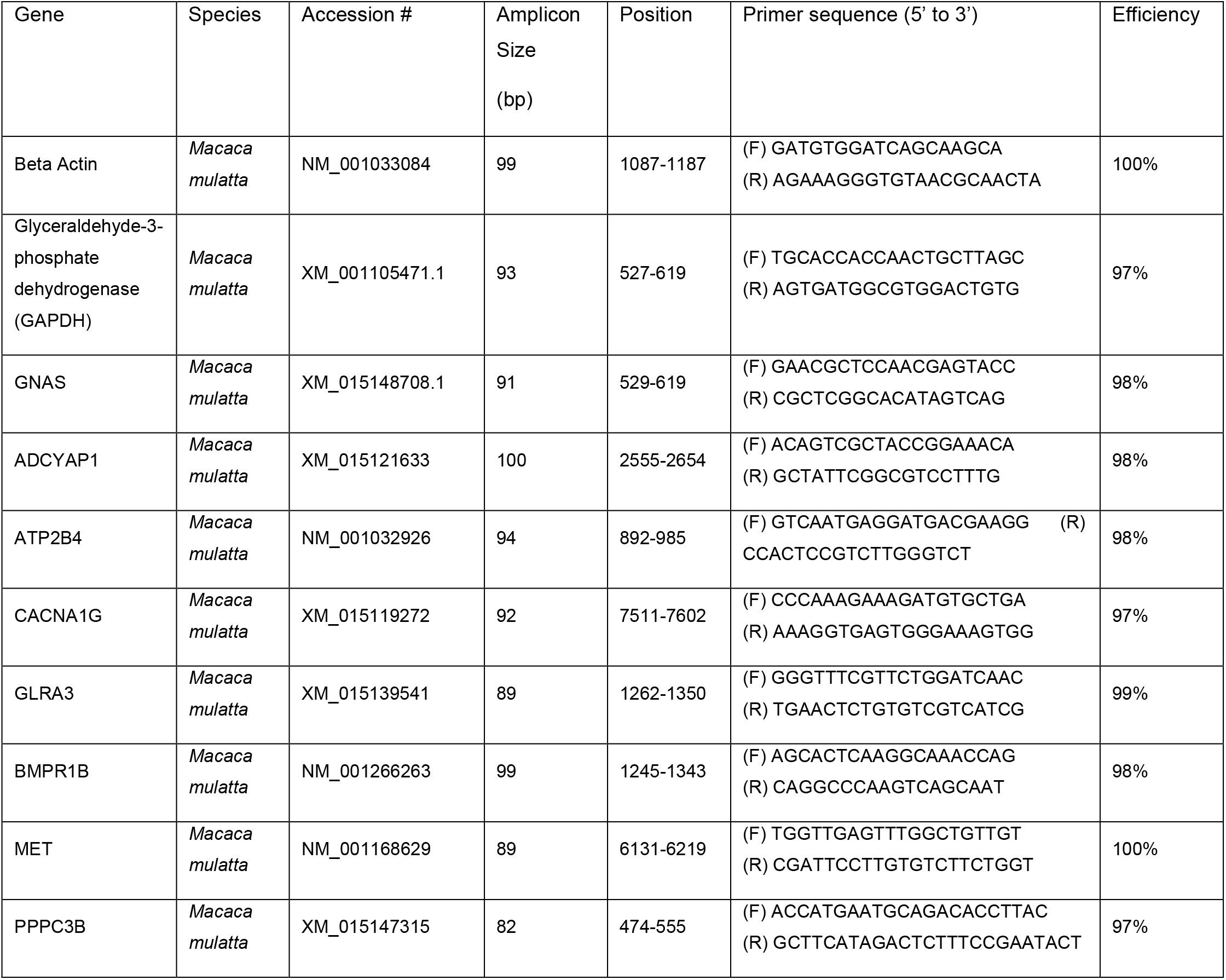
Oligonucleotide primers used in quantitative PCR

**Table 2.** Sholl Analysis of apical and basal dendrites

| Variable | DLPFC L3PNs<br>(n=21) | PPC L3PNs<br>(n=18) | Student's t-test | p value |
| --- | --- | --- | --- | --- |
| Peak number of intersections,<br>Apical oblique branches | 16.8 ± 0.9 | 19.4 ± 1.7 | $t_{(37)} = 1.422$ | 0.1634 |
| Peak number of intersections<br>Apical tuft | 12.0 ± 1.1 | 12.2 ± 1.5 | $t_{(37)} = 0.093$ | 0.9262 |
| Peak number of intersections<br>Basal dendrites | 33.1 ± 1.9 | 22.0 ± 1.8 | $t_{(37)} = 4.158$ | 0.000183 |
| Peak length (µm)<br>Apical oblique branches | 404 ± 23 | 475 ± 53 | $t_{(37)} = -1.290$ | 0.2050 |
| Peak length (µm)<br>Apical tuft | 346 ± 43 | 291 ± 39 | $t_{(37)} = 0.929$ | 0.356 |
| Peak length (µm)<br>Basal dendrite | 1058 ± 84 | 590 ± 56 | $t_{(37)} = 4.467$ | 0.000072 |

In contrast to the absence of difference in L3PN apical dendrites between areas, the basal dendrites were significantly larger in DLPFC L3PNs than PPC L3PNs (Figure 4E), as reflected in both the total length (~54 % larger) and convex hull volume (~43% larger). Moreover, Sholl analysis showed robust differences in basal dendrite length (Figure 4F) and complexity (Figure 4G) between L3PNs of DLPFC and PPC. These differences were confirmed by analysis of scaled Sholl plots (Figure 4H), and by comparisons of the peak length and peak number of intersections (Table 2). Importantly, the number of primary basal dendrites originating from the soma was similar between DLPFC (6.4 ± 0.5 dendrites, n=21) and PPC L3PNs (5.9 ± 0.3 dendrites, n=18; p=0.296, Mann Whitney U test). Hence, the greater length and complexity of DLPFC versus PPC L3PN basal dendrites were associated with different properties of individual basal dendritic trees.

Our data suggest that, at the single-cell level, basal dendrites contain a larger fraction of the total dendrite length in DLPFC L3PNs compared to PPC L3PNs. Hence, we computed the ratio between basal dendrite and total dendrite length for individual L3PNs, and found that this ratio was significantly greater in DLPFC L3PNs (0.475 ± 0.019, n=21) than PPC L3PNs (0.380 ± 0.017, n=18; t(37)= 3.775, p=0.00056). Similar findings were obtained for the convex hull volume values (data not shown).

The larger and more complex basal dendritic trees of DLPFC L3PNs are consistent with integration of larger numbers of synaptic inputs. However, the number of inputs integrated also depends on input density. Thus, we estimated excitatory input density in basal dendrites by measuring the density of dendritic spines (Figure 5A,B). As in previous studies of L3PNs from monkey neocortex (Elston and Rosa, 1997; Elston et al., 1999; Elston et al., 2011a; Medalla and Luebke, 2015; Gilman et al., 2017), spine density was low in the proximal basal dendrites, and increased with distance from the soma to reach a plateau by mid dendrite before declining near the distal end (Figure 5C). Despite similar spatial profiles in DLPFC and PPC L3PNs, spine density was higher in DLPFC L3PNs across multiple compartments (Figure 5C,D). Furthermore, both the peak spine density (Figure 5E) and the mean spine density (DLPFC L3PNs, n=20, 0.954 ± 0.036 spines/μm; PPC L3PNs, n=15, 0.673 ± 0.049 spines/μm; t_(33)_=4.691, p=0.000046) were higher in DLPFC L3PN basal dendrites. The total number of basal dendrite spines per L3PN, estimated using the mean spine density and total basal dendrite length, was 89% higher in DLPFC L3PNs (DLPFC: 5082 ± 530 spines; PPC: 2689 ± 365 spines, t_(28)_=3.609, p=0.0012).

**Figure 5.**
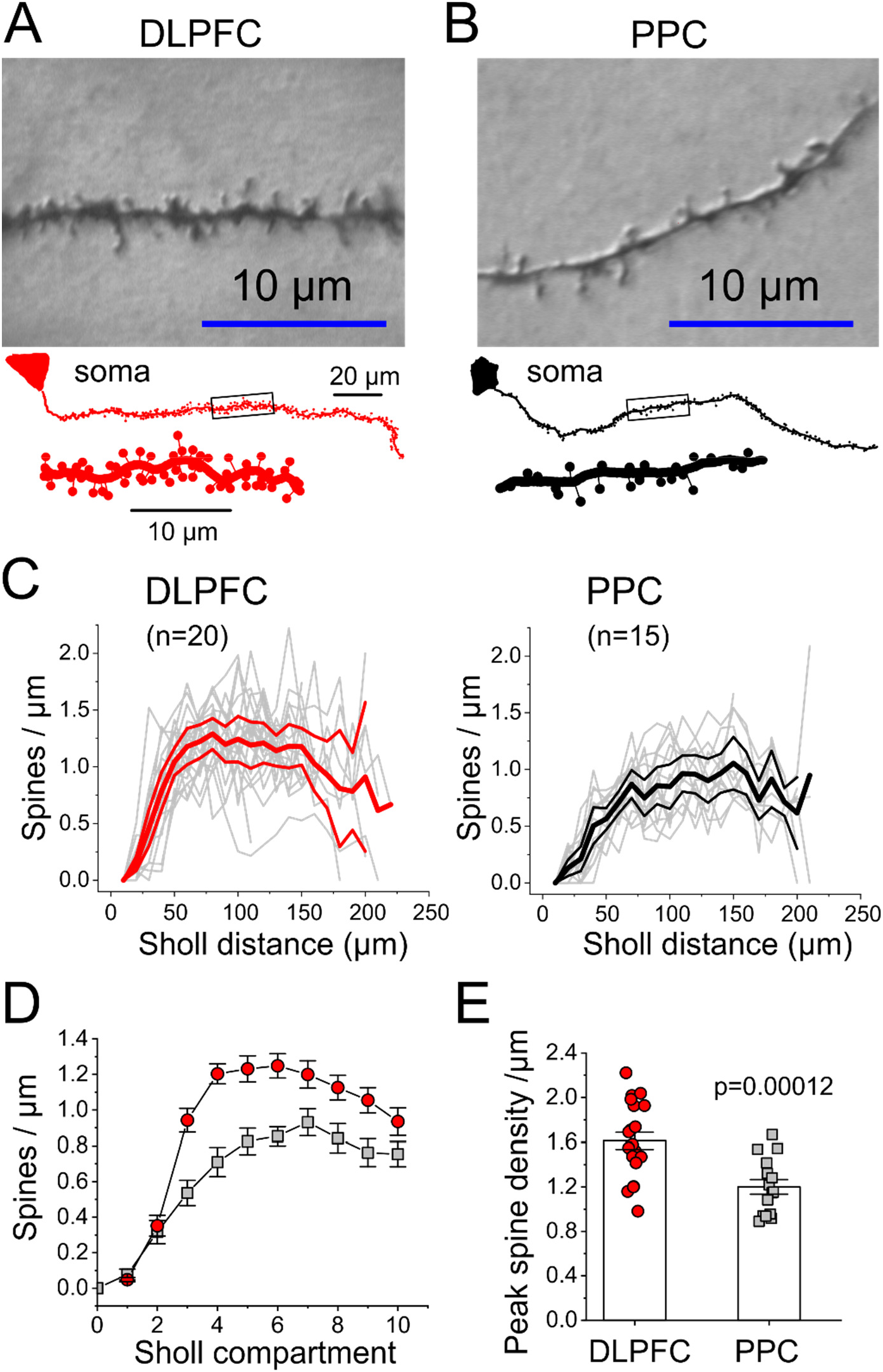
Quantitative analysis of dendritic spine density in basal dendrites of DLPFC and PPC L3PNs. **A)** Differential interference contrast image of a segment of mid basal dendrite from a L3PN from DLPFC, showing multiple dendritic spines with various spine morphologies. The bottom panel shows the reconstruction of a basal dendrite and its dendritic spines between soma and distal dendrite end. The dendrite segment identified by the box is shown in a zoomed-in view below the dendrite. **B)** Image and reconstructions as in A), but for a L3PN from PPC. **C)** Sholl plots of dendritic spine density for the basal dendrites of DLPFC (n=20) and PPC (n=15) L3PNs. The thick lines indicate the mean value and the dark lines above and below are the 95% confidence intervals. The gray lines represent data from individual L3PNs. **D)** Scaled Sholl plots of dendritic spine density in the basal dendrites of the DLPFC and PPC L3PNs illustrated in C). The basal dendrites of each L3PN were divided into 10 compartments. **E)** Bar graphs of the peak spine density in the basal dendrites of the DLPFC and PPC L3PNs illustrated in C). Shown are the results of a Student’s t-test comparison.

### The differences in basal dendrite properties between cortical areas are independent of physiological subtype

The differences in basal dendrites between DLPFC and PPC L3PNs could be attributed to the larger percentage of B-L3PNs in DLPFC than PPC (Figure 1E), if, for instance, B-L3PNs have larger and more complex dendrites. Thus, we compared the dendritic tree properties of the RS-L3PNs and B-L3PNs that had dendritic tree reconstructed. In this sample of L3PNs, the proportions of RS-L3PNs to B-L3PNs in DLPFC (11:10) and in PPC (17:1), were very similar to those observed in the total sample (Figure 1E). PPC B-L3PNs were excluded from the statistical analysis, because only a single B-L3PN from PPC had dendrites reconstructed. Comparisons of apical dendrite properties between the other three groups (data not shown) did not reveal significant differences within or between areas (Length: F_(2,35)_=0.0064, p=0.994; Convex hull volume: F_(2,35)_=0.317, p=0.730). Contrasting with the absence of difference in apical dendrites, the basal dendrite length was larger in RS-L3PNs from DLPFC than PPC (Figure 6A) but did not differ between RS-L3PNs and B-L3PNs within DLPFC (Figure 6A). The basal dendrites of RS-L3PNs had larger convex hull volume in DLPFC than PPC (Figure 6B). However, this difference was not significant (F_(2, 35)_=2.437, p=0.102).

**Figure 6.**
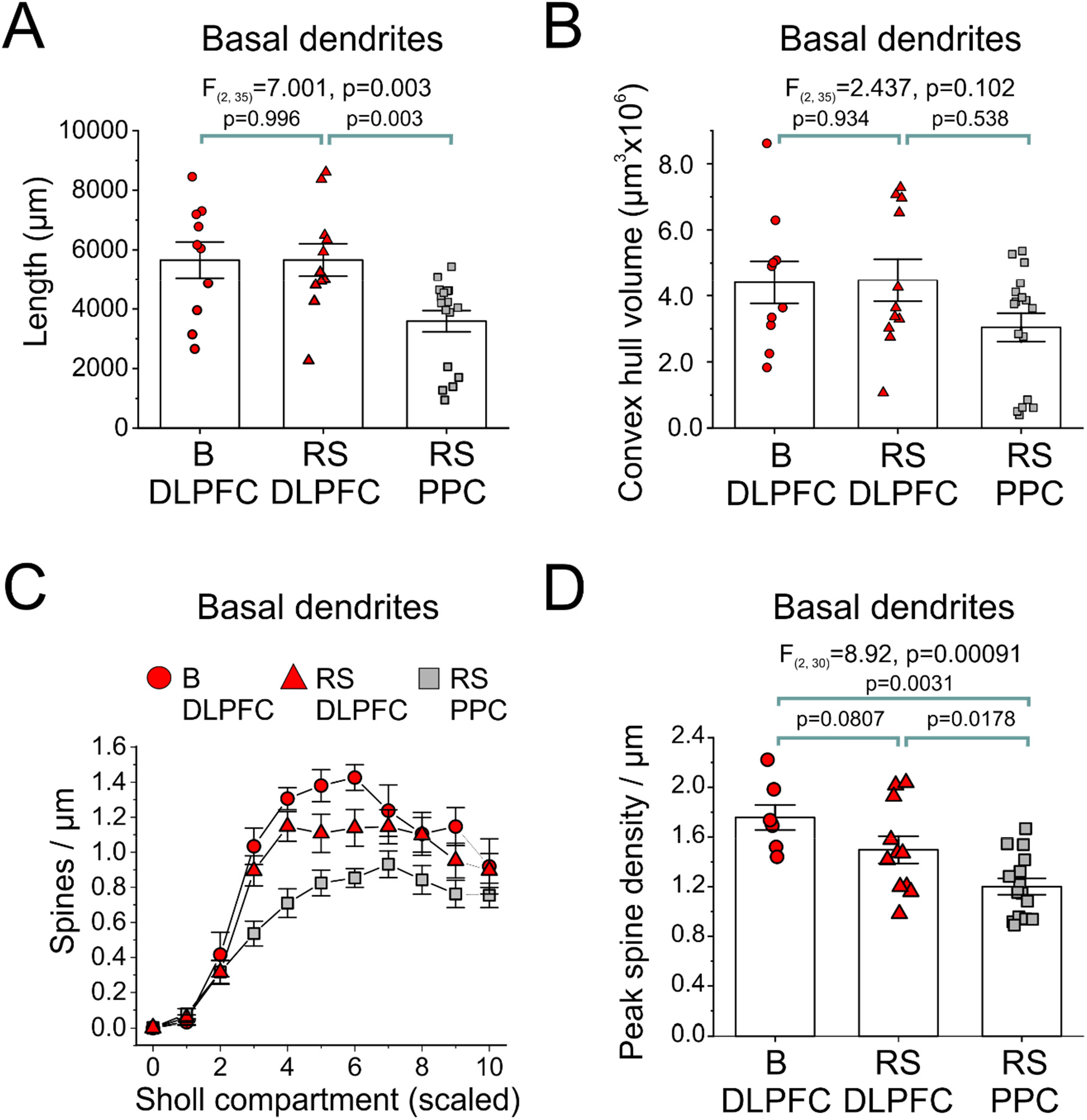
Quantitative analysis of basal dendrite properties and spine density in DLPFC and PPC L3PNs identified as RS-L3PNs and B-L3PNs. **A)** Bar graphs of total length of basal dendrites for B-L3PNs (here abbreviated as B) and RS-L3PNs (here abbreviated as RS) from DLPFC and PPC. Shown are the results of One-Way ANOVA followed by post-hoc comparisons. Note that B-L3PNs from PPC were not included in the graphs or the statistical comparisons because only a single B-L3PNs had dendrites reconstructed. **B)** Bar graphs of convex hull volume for B and RS L3PNs from DLPFC and PPC. Results of statistical analysis reported as in A). **C)** Scaled Sholl plots of dendritic spine density for B-L3PNs from DLPFC and RS-L3PNs from DLPFC or PPC. The reconstructed basal dendrite of each L3PN was divided into 10 compartments. **D)** Bar graphs of peak dendritic spine density in basal dendrites for B and RS L3PNs from DLPFC and PPC. Shown are the results of One-Way ANOVA followed by post-hoc comparisons. Note that B-L3PNs from PPC were not included in the graphs or the statistical comparisons because only a single B-L3PNs had dendrites reconstructed.

Next, we investigated if the higher spine density in DLPFC L3PN basal dendrites could be attributed to higher spine density in B-L3PNs. We found that spine density was higher across various compartments of the basal dendrites of DLPFC L3PNs (Figure 6C). Moreover, PPC RS-L3PNs had significantly lower basal dendrite spine density than either RS-L3PNs or B-L3PNs from DLPFC, whereas the peak spine density was highest in DLPFC B-L3PNs (Figure 6D). One-Way ANOVA analysis revealed that the total number of basal dendrite spines differed significantly between L3PN groups (One-Way ANOVA F_(2,30)_ = 7.798, p=0.0019). Total spine numbers did not differ between DLPFC B-L3PNs (5343 ± 934 spines) and DLPFC RS-L3PNs (5065 ± 536 spines, p=0.746), but the spine number in PPC RS-L3PNs (2746 ± 345 spines) was lower than those in either B-L3PNs (p=0.0031) or RS-L3PNs (p=0.0024) from DLPFC. Hence, the differences in basal dendrite length, complexity, and spine density between DLPFC and PPC L3PNs cannot be attributed to the higher proportion of B-L3PNs in DLPFC.

### DLPFC and PPC L3PNs differ in gene expression profiles

To determine if the observed differences in physiological and morphological properties reflect regional differences in L3PN gene expression, we combined laser microdissection with microarray profiling to compare the transcriptomes of DLPFC and PPC L3PNs. Using tissue from 5 of the animals studied in the ex vivo (brain slice) experiments, we found that 753 probes representing 678 unique transcripts were differentially expressed (q<0.1) in L3PNs between DLPFC and PPC as illustrated in Figure 7A. Of the 678 unique transcripts, 636 were at least partially annotated. Of the 636 transcripts, approximately half (n=315) had higher expression levels in DLPFC than PPC, and the remainder (n=321) had higher expression in PPC. Table 3 reports the top 20 genes found to be differentially expressed by L3PNs in DLPFC area 46 or PPC areas 7a and LIP.

**Figure 7.**
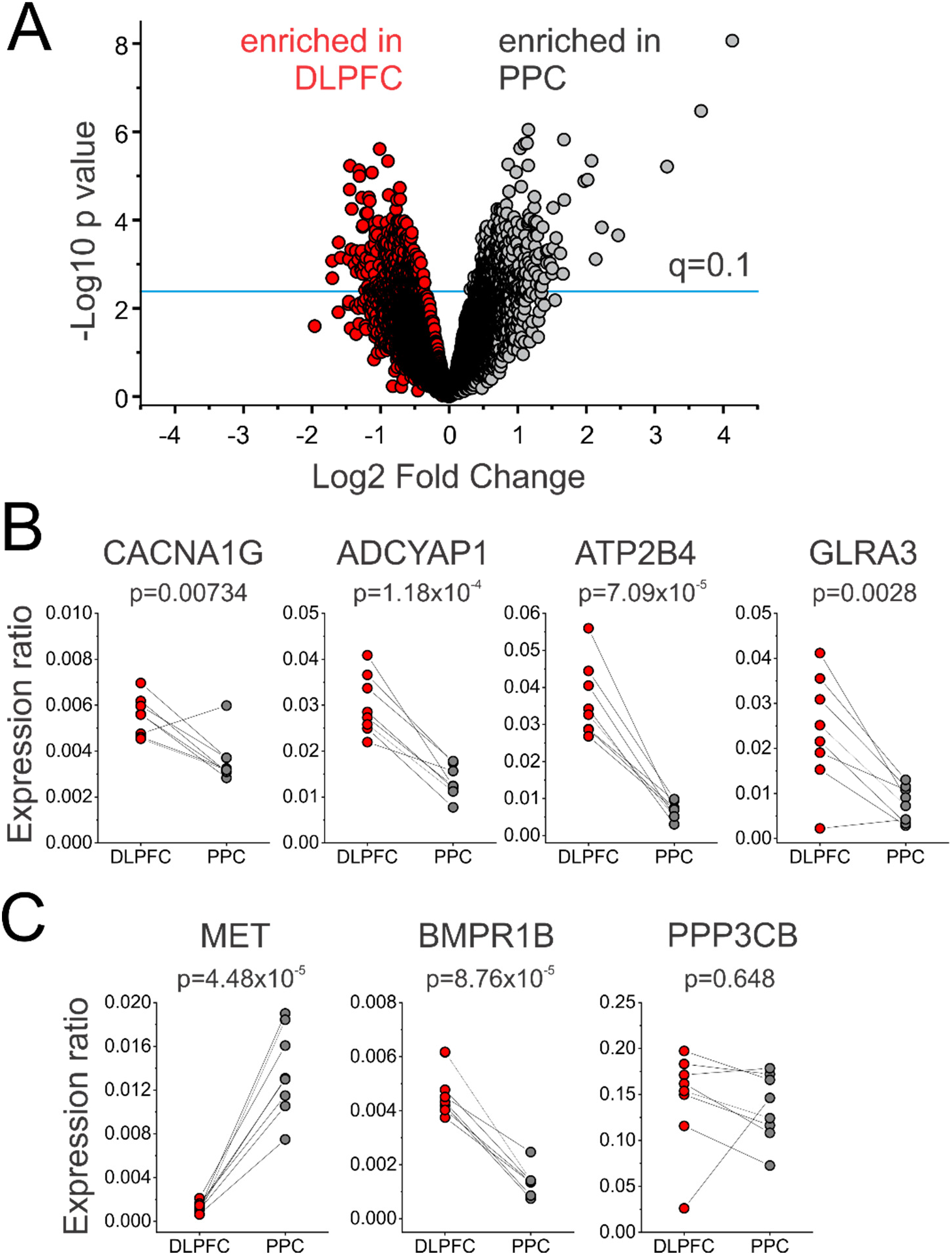
Profiling of layer 3 pyramidal neurons from DLPFC and PPC. **A)** Volcano plot of all probes following microarray profiling. The horizontal line represents the statistical cutoff at q=0.1. **B)** qPCR validation for 4 transcripts enriched in DLPFC. The lines join data from DLPFC and PPC samples of an individual animal. Shown are the results of Student’s t-test comparisons. **C)** qPCR validation for 3 transcripts enriched in PPC. The lines join data from DLPFC and PPC samples of an individual animal. Shown are the results of Student’s t-test comparisons.

**Table 3.**
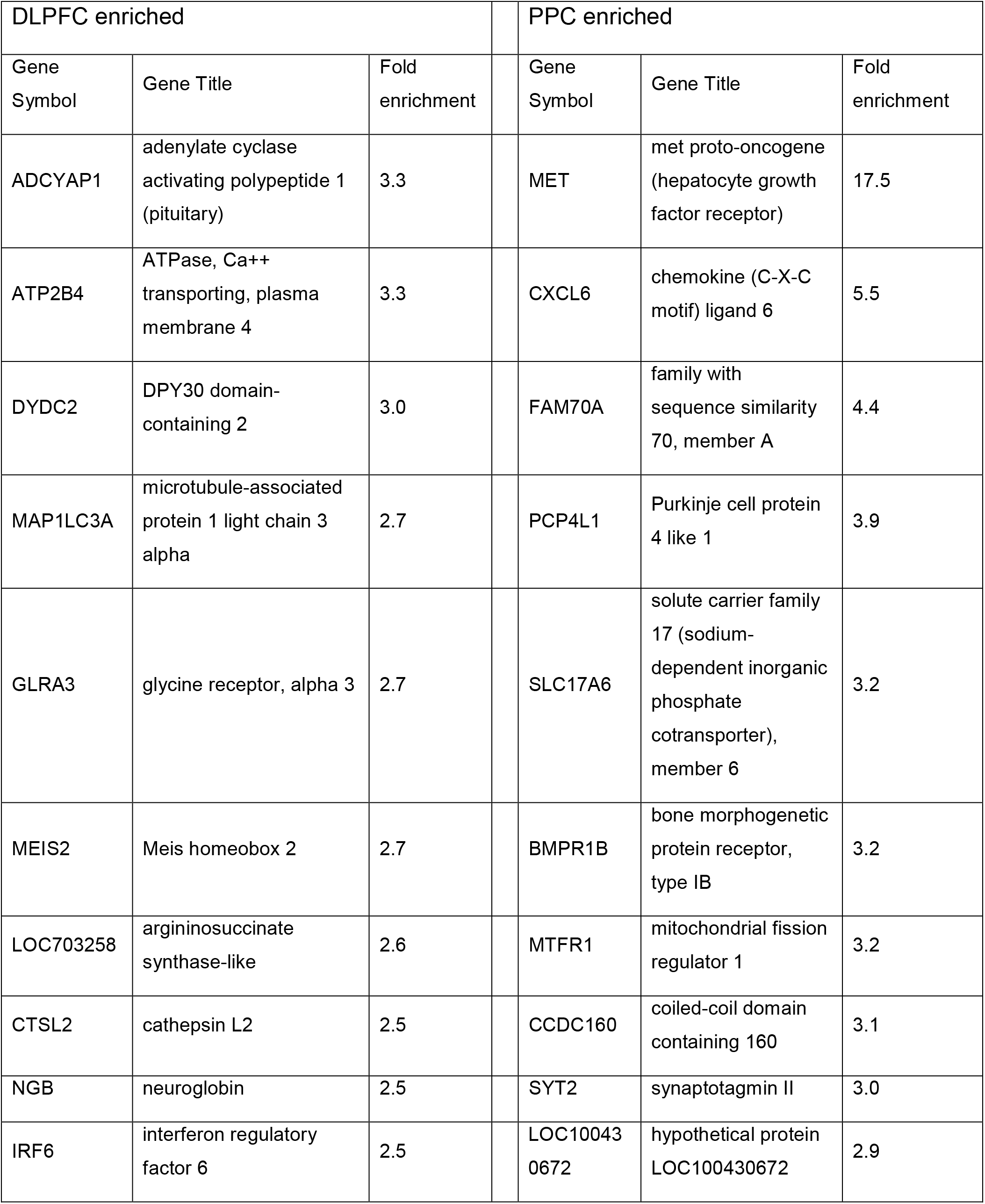
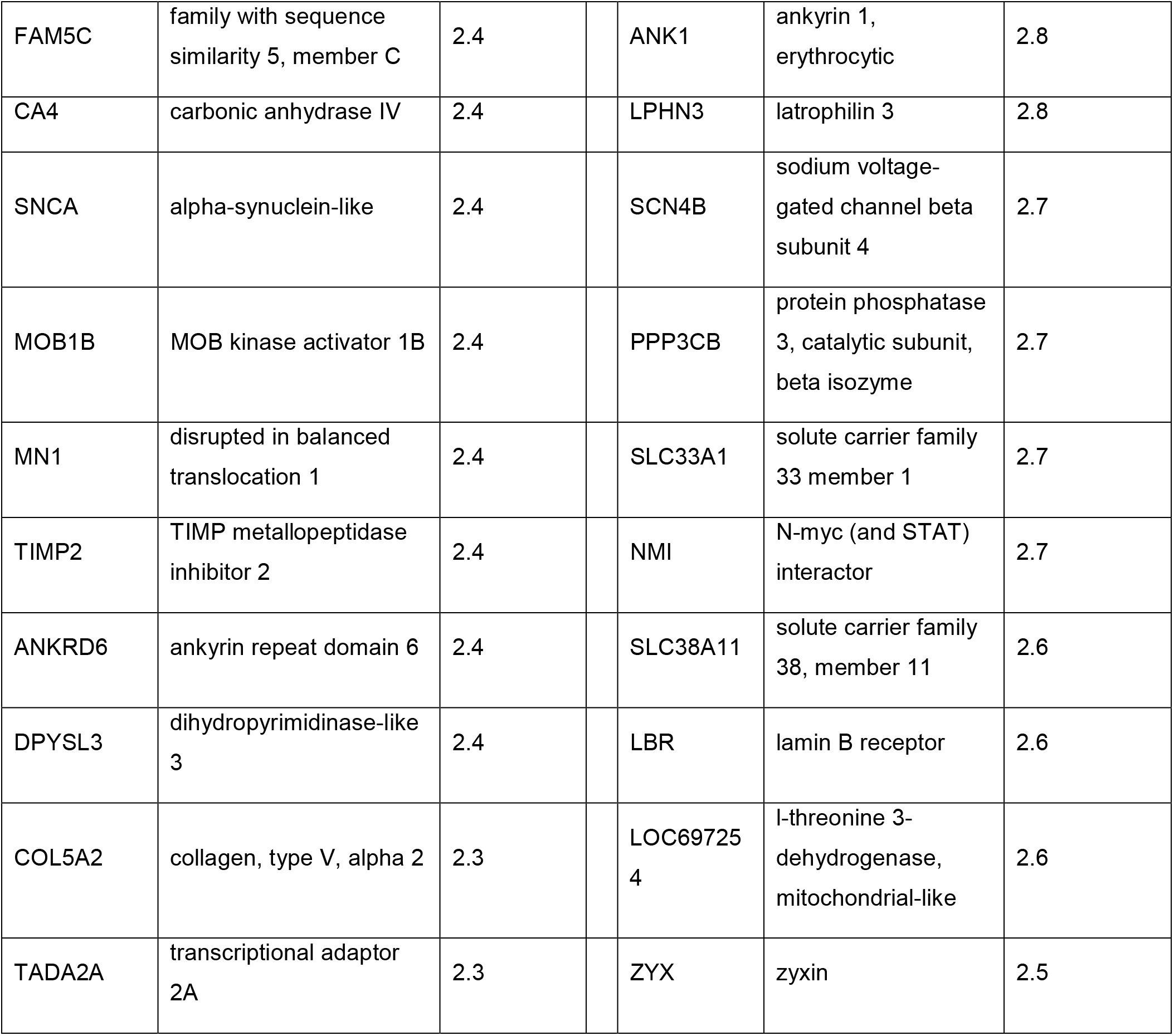
Top 20 genes differentially expressed by L3PNs from DLPFC and PPC

| DLPFC enriched |  |  | PPC enriched |  |  |
| --- | --- | --- | --- | --- | --- |
| Gene Symbol | Gene Title | Fold enrichment | Gene Symbol | Gene Title | Fold enrichment |
| ADCYAP1 | adenylate cyclase activating polypeptide 1 (pituitary) | 3.3 | MET | met proto-oncogene (hepatocyte growth factor receptor) | 17.5 |
| ATP2B4 | ATPase, Ca<sup>++</sup> transporting, plasma membrane 4 | 3.3 | CXCL6 | chemokine (C-X-C motif) ligand 6 | 5.5 |
| DYDC2 | DPY30 domain-containing 2 | 3.0 | FAM70A | family with sequence similarity 70, member A | 4.4 |
| MAP1LC3A | microtubule-associated protein 1 light chain 3 alpha | 2.7 | PCP4L1 | Purkinje cell protein 4 like 1 | 3.9 |
| GLRA3 | glycine receptor, alpha 3 | 2.7 | SLC17A6 | solute carrier family 17 (sodium-dependent inorganic phosphate cotransporter), member 6 | 3.2 |
| MEIS2 | Meis homeobox 2 | 2.7 | BMPR1B | bone morphogenetic protein receptor, type IB | 3.2 |
| LOC703258 | argininosuccinate synthase-like | 2.6 | MTFR1 | mitochondrial fission regulator 1 | 3.2 |
| CTSL2 | cathepsin L2 | 2.5 | CCDC160 | coiled-coil domain containing 160 | 3.1 |
| NGB | neuroglobin | 2.5 | SYT2 | synaptotagmin II | 3.0 |
| IRF6 | interferon regulatory factor 6 | 2.5 | LOC100430672 | hypothetical protein LOC100430672 | 2.9 |

|  |  |  |  |  |  |  |
| --- | --- | --- | --- | --- | --- | --- |
| FAM5C | family with sequence similarity 5, member C | 2.4 |  | ANK1 | ankyrin 1, erythrocytic | 2.8 |
| CA4 | carbonic anhydrase IV | 2.4 |  | LPHN3 | latrophilin 3 | 2.8 |
| SNCA | alpha-synuclein-like | 2.4 |  | SCN4B | sodium voltage-gated channel beta subunit 4 | 2.7 |
| MOB1B | MOB kinase activator 1B | 2.4 |  | PPP3CB | protein phosphatase 3, catalytic subunit, beta isozyme | 2.7 |
| MN1 | disrupted in balanced translocation 1 | 2.4 |  | SLC33A1 | solute carrier family 33 member 1 | 2.7 |
| TIMP2 | TIMP metalloproteinase inhibitor 2 | 2.4 |  | NMI | N-myc (and STAT) interactor | 2.7 |
| ANKRD6 | ankyrin repeat domain 6 | 2.4 |  | SLC38A11 | solute carrier family 38, member 11 | 2.6 |
| DPYSL3 | dihydropyrimidinase-like 3 | 2.4 |  | LBR | lamin B receptor | 2.6 |
| COL5A2 | collagen, type V, alpha 2 | 2.3 |  | LOC697254 | l-threonine 3-dehydrogenase, mitochondrial-like | 2.6 |
| TADA2A | transcriptional adaptor 2A | 2.3 |  | ZYX | zyxin | 2.5 |

Some of the genes identified near the top of the rank of differential expression in the microarray analysis (Table 3) might contribute to the regional differences in L3PN phenotype. For example, *MET* (17.5 fold higher in PPC L3PNs), encodes a tyrosine kinase receptor involved in control of dendritic arbor properties and spine density (Eagleson et al., 2017). Other genes near the top of the rank of differential expression (Table 3), such as *ADCYAP1* (3.3 fold higher in DLPFC L3PNs) and *GLRA3* (2.7 fold higher in DLPFC L3PNs), are markers of specific projection subtypes of PNs (Sorensen et al., 2015; Chevee et al., 2018). Several other transcripts enriched in DLPFC L3PNs (*ATP2B4*, 3.3 fold), or in PPC L3PNs (*PPP3CB*, 2.7 fold; *BMPR1B*, 3.2 fold), encode proteins involved in cellular metabolism (*ATP2B4, PPP3CB*) or in dendritic morphogenesis (*BMPR1B*).

Genes for voltage-gated Na^+^, K^+^ and Ca^2+^ channels were expressed in significant levels by both DLPFC and PPC L3PNs. However, in accordance with the modest regional differences in single-cell excitability, for most of these ion channel genes the differences in expression levels between DLPFC and PPC L3PNs were not significant (data not shown). Interestingly, among genes identified as differentially expressed in our microarray analysis is *CACNA1G* (1.9 fold higher in DLPFC L3PNs). *CACNA1G* encodes a subunit of the T-type voltage-gated calcium channel family Ca_v_3, involved in the generation of action potential bursts in PNs (Williams and Stuart, 1999; Clarkson et al., 2017; Dumenieu et al., 2018). Consistent with the well-known body of data from rodent cortex showing that burst-firing PNs are widely distributed across cortical regions, *CACNA1G* is expressed in multiple regions of the monkey neocortex (Bernard et al., 2012). In agreement with our finding that the proportion of bursting L3PNs differs between DLPFC and PPC, *CACNA1G* expression levels differ significantly across areas of the monkey neocortex (Bernard et al., 2012).

To confirm the differential regional expression of transcripts revealed by the microarray data, we captured L3PNs in a separate cohort of monkeys (see Materials and Methods) and used qPCR to quantify the expression levels of some of the genes that might underlie the phenotypic differences between DLPFC and PPC L3PNs. For transcripts identified by the microarray profiling as DLPFC-enriched, all of those assessed by qPCR (*CACNA1G, ADCYAP1, GLRA3* and *ATP2B4*), were confirmed to display higher expression in L3PNs from DLPFC than PPC (Figure 7B). Furthermore, among the transcripts identified by microarray profiling as PPC-enriched, 2 of the 3 transcripts assessed with qPCR (*MET* and *BMPR1B*, but not *PPP3CB*) also showed significantly higher expression in L3PNs from PPC than DLPFC (Figure 7C).

The differential gene expression between DLPFC and PPC L3PNs revealed by our microarray data and confirmed by qPCR, is in addition consistent with data from a study of the transcriptional architecture of the macaque monkey cortex (Bernard et al., 2012). For instance, several of the genes highlighted in Table 3 were previously shown to differ in expression levels across areas of the monkey cortex (Bernard et al., 2012). These include 13 genes with higher expression in DLPFC versus PPC L3PNs (*ATP2B4, DYDC2, MAP1LC3A, GLRA3, MEIS2, NGB, IRF6, FAM5C, CA4, SNCA, TIMP2, ANKRD6*, and *TADA2A*), as well as 12 genes with expression enriched in PPC L3PNs versus DLPFC L3PNs (*MET, FAM70A, PCP4L1, BMPR1B, CCDC160, ANK1, PPP3CB, NMI, SLC38A11, LBR, LOC697254*, and *ZYX*).

For some genes here identified as differentially expressed in L3PNs from DLPFC versus PPC (Table 3), expression levels did not differ across areas of the monkey cortex in a prior study (Bernard et al., 2012). This group includes 7 genes from Table 3 with higher expression levels in DLPFC L3PNs (*ADCYAP1, LOC703258, CTSL2, MOB1B, MN1, DPYSL3*, and *COL5A2*), and 8 genes enriched in PPC L3PNs (*CXCL6, SLC17A6, MTFR1, SYT2, LOC100430672, LPHN3, SCN4B*, and *SLC33A1*). However, unlike our study capturing L3PN cell bodies, Bernard et al. 2012 studied the entire cortical layer 3. Thus, the differences between our findings and those of the previous study (Bernard et al., 2012) may reflect transcripts that are differentially expressed across regions in a cell type-specific manner.

## Discussion

We found that L3PNs of monkey DLPFC and PPC could be classified according to two major firing pattern subtypes. DLPFC contained equal proportions of regular spiking and bursting L3PNs, whereas in PPC ~95% of the L3PNs were of the regular spiking subtype. Morphological reconstructions showed that, relative to PPC L3PNs, DLPFC L3PNs have larger and more complex basal dendrites with higher spine density. Finally, significant differences in gene expression revealed by transcriptome analysis suggest a molecular basis for the differences in L3PN phenotypes between areas.

### Physiological properties of L3PNs in DLPFC and PPC

The timescale of fluctuations in single-neuron activity changes gradually across the monkey neocortex, with sensory cortices and DLPFC exhibiting the longest and shortest timescales, respectively (Murray et al., 2014). A shorter timescale might be associated with a shorter neuronal membrane time constant. However, although PPC neurons exhibit an activity timescale intermediate between sensory cortex and DLPFC (Murray et al., 2014), we did not find differences in the membrane time constant between DLPFC and PPC L3PNs, nor did the time constant differ between V1 and DLPFC L3PNs (Amatrudo et al., 2012). Hence, the variation in activity timescales across primate cortex might not be associated with differences in the biophysical properties underlying temporal integration by L3PNs.

L3PNs from monkey V1 are intrinsically more excitable than DLPFC L3PNs (Amatrudo et al., 2012; Gilman et al., 2017). As V1 L3PNs also have smaller dendrites with lower spine density, they may receive lower levels of excitatory drive which, via homeostatic mechanisms, may induce greater excitability (Spruston, 2008; Debanne et al., 2019). Despite differences in dendritic tree size and spine density, however, we found very modest differences in single-cell excitability between DLPFC and PPC L3PNs. Future studies are thus needed to assess the levels of excitatory drive in DLPFC and PPC L3PNs, and whether the intrinsic excitability of L3PNs is homeostatically adjusted by the excitatory drive in these cortical areas.

### Dendritic tree properties and spine density in DLPFC and PPC L3PNs: comparison with previous studies

Our estimates of apical and basal dendritic tree parameters are highly consistent with previous studies of L3PNs filled with biocytin in monkey DLPFC slices (Amatrudo et al., 2012; Gonzalez-Burgos et al., 2015; Medalla and Luebke, 2015; Gilman et al., 2017). Most studies of L3PN dendrites in primate neocortex, however, were performed in aldehyde-fixed tissue, and assessed only basal dendrites (Elston and Fujita, 2014). Our estimates of dendritic size parameters are generally larger than those for L3PNs visualized in fixed tissue from either DLPFC (Anderson et al., 1995; Soloway et al., 2002; Duan et al., 2003; Kabaso et al., 2009) or PPC (Motley et al., 2018).

In addition, the basal dendrite spine density estimated here is consistent with previous studies of DLPFC L3PNs biocytin-filled in slices (Amatrudo et al., 2012; Gilman et al., 2017) or PPC L3PNs microinjected in fixed tissue (Motley et al., 2018), but is higher than estimates in most studies in fixed tissue (Anderson et al., 1995; Elston and Rosa, 1997, 2000; Kabaso et al., 2009; Elston et al., 2011b; Young et al., 2014). Notably, previous studies of dendritic tree properties in aldehyde-fixed tissue found differences between L3PNs from PPC area 7a and orbitofrontal areas 10, 11 or 12 in monkeys (Elston, 2000; Elston et al., 2001). Thus, despite discrepancy between methods, possibly due to differences in tissue shrinkage or the fraction of total dendritic tree reconstructed, experiments in brain slices and fixed tissue are consistent in revealing regional differences in L3PN dendritic tree properties across areas of the primate cortex.

### Gene expression profiles of L3PNs in DLPFC and PPC

Transcriptome analysis revealed that DLPFC and PPC L3PNs differentially expressed hundreds of genes, for many of which expression levels were previously found to vary across regions of the monkey neocortex (Bernard et al., 2012). However, other genes differentially expressed by DLPFC versus PPC L3PNs may distinguish these cell types, as they were not differentially expressed between areas of monkey cortex when whole cortical layers were analyzed (Bernard et al., 2012).

Several of the genes differentially expressed may contribute to the different phenotypes of DLPFC and PPC L3PNs. Among these is *MET*, a gene strongly associated with risk for autism (Eagleson et al., 2017), that encodes the MET receptor strongly expressed by PNs (Kast et al., 2017). Reducing *MET* expression increases L3PN basal dendrite size and complexity (Judson et al., 2010). Therefore, higher *MET* levels in PPC L3PNs are consistent with smaller dendritic tree length and complexity. MET, however, also plays roles unrelated to regulating dendrite morphology (Xie et al., 2016), and its regulation of dendrite properties varies among cortical layers and cell types, and between basal and apical dendrites (Judson et al., 2010; Heun-Johnson and Levitt, 2018). Thus, further studies are needed to assess cause-effect relations between MET levels and dendritic tree properties in DLPFC and PPC L3PNs.

The main electrophysiological difference observed between DLPFC and PPC L3PNs was a higher proportion of bursting L3PNs in DLPFC. Burst firing depends on voltage-gated Ca^2+^ channels (Williams and Stuart, 1999; Clarkson et al., 2017; Dumenieu et al., 2018) of the T-type Ca_v_3 family (Nanou and Catterall, 2018). Therefore, our finding that expression of the Ca_v_3 alpha 1G subunit gene *CACNA1G* was enriched in DLPFC L3PN samples, is consistent with the larger proportion of B-L3PNs in DLPFC.

### Functional relevance of the differences between DLPFC and PPC L3PNs

We found that DLPFC L3PNs have greater basal dendrite length, complexity, and spine number. The functional properties of basal dendrites remain unexplored in primate cortex, but in rodent cortex basal dendrites display NMDA spikes (Schiller et al., 2000) that counteract signal attenuation (Nevian et al., 2007). NMDA receptor activation may contribute to recurrent excitation (Lisman et al., 1998), and thus to persistent firing and gamma oscillations, the activity patterns that might mediate working memory (Constantinidis et al., 2018; Lundqvist et al., 2018). Recurrent excitation primarily involves inputs onto basal dendrite spines (Markram et al., 1997; Gökçe et al., 2016), and NMDA spikes are evoked most efficiently by stimulating clusters of axospinous synapses within basal dendrites (Polsky et al., 2004). Thus, by facilitating NMDA spike production, the higher spine density in DLPFC L3PNs may be crucial for working memory-related activity.

The main physiological distinction between cortical areas was a larger fraction of burst-firing L3PNs in DLPFC. By signaling the slope of low frequency fluctuations in excitatory input (Kepecs et al., 2002), burst firing may enhance transmission of alpha band rhythms thought to convey working memory-related top-down signals (Miller et al., 2018; Quentin et al., 2019). Thus, DLPFC B-L3PNs may specialize in transmitting top-down signals to other cortical regions. Burst firing might also enhance information transmission by spike trains in neuronal ensembles, enabling PN groups to simultaneously process of top-down and bottom-up streams of information (Naud and Sprekeler, 2018). Thus, the higher proportion of burst-firing L3PNs observed here in DLPFC might equip DLPFC neuronal ensembles with a richer repertoire of information-processing mechanisms. Interestingly, in agreement with our data in brain slices, nearly half of the neurons (putative PNs) recorded from monkey DLPFC during behavioral tasks are burst-firing (Ardid et al., 2015; Voloh and Womelsdorf, 2018). It is unclear, however, if burst-firing cells in the DLPFC of behaving monkeys are L3PNs, or cells in layer 5, where bursting PNs are more abundant (Chang and Luebke, 2007).

Spike frequency adaptation destabilizes persistent firing (Carter and Wang, 2007), an activity pattern possibly mediating working memory storage (Constantinidis et al., 2018), however see (Lundqvist et al., 2018). As RS-L3PNs had weaker spike frequency adaptation than B-L3PNs, one possibility is that the RS phenotype facilitates persistent firing. Hence, in DLPFC neuron ensembles, B-L3PNs may transmit top-down information for processes such as cognitive control, and RS-L3PNs may mediate storage and/or maintenance of task-relevant rules. Conversely, the predominantly RS-L3PN PPC population may be optimized for storage of items in working memory, as suggested by neuroimaging studies (Hahn et al., 2018). These studies showed that storage capacity is strongly correlated with the PPC BOLD signal, and that, in schizophrenia, such correlation is disrupted in association with decreased PPC activation and storage capacity deficits (Hahn et al., 2018).

Our findings of differential expression of hundreds of genes suggest that the phenotypic differences between DLPFC and PPC L3PNs have a molecular basis. However, further evaluation by single-cell transcriptomics and gene pathway/transcriptional network analysis is needed to fully characterize the differential gene expression underlying the differences between areas, and to identify molecular subtypes of L3PNs within monkey DLPFC and PPC. As the layer 3 neuron transcriptomes in macaque and human cortices are closely related (Zhu et al., 2018), our data assessing monkey L3PN properties will help identifying how gene expression distinguishes types of L3PNs in the human cortex. Additionally, our data may facilitate understanding the causes and consequences of altered gene expression in schizophrenia, a disease that affects the neuronal transcriptome in the DLPFC, the PPC, and other areas of the working memory network (Arion et al., 2015; Hoftman et al., 2018; Tsubomoto et al., 2018).

## Acknowledgements

We thank Dr. Tatiana Tikhonova for help in the initial stage of this project; Sam Dienel and Liban Dinka for help with laser microdissection; and Kate Gurnsey for assistance in surgical procedures.

